# AAV-HBV mouse model replicates immune exhaustion patterns of chronic HBV patients at single-cell level

**DOI:** 10.1101/2023.08.07.552328

**Authors:** Nádia Conceição-Neto, Qinglin Han, Zhiyuan Yao, Wim Pierson, Qun Wu, Koen Dockx, Liese Aerts, Dries De Maeyer, Koen Van den Berge, Chris Li, George Kukolj, Ren Zhu, Ondřej Podlaha, Isabel Nájera, Ellen Van Gulck

## Abstract

**Background and Aims:** Unresolved hepatitis B virus (HBV) infection leads to a progressive state of immune exhaustion that impairs resolution of infection, leading to chronic infection (CHB). The immune-competent AAV-HBV mouse is a common HBV preclinical immune competent model, though a comprehensive characterization of the liver immune microenvironment and its translatability to human infection is still lacking. We investigated the intrahepatic immune profile of the AAV-HBV mouse model at a single-cell level and compared with data from CHB patients in immune tolerant (IT) and immune active (IA) clinical stages.

**Methods:** Immune exhaustion was profiled through an iterative subclustering approach for cell-typing analyses of single-cell RNA-sequencing data in CHB donors and compared to the AAV-HBV mouse model 24-weeks post-transduction to assess its translatability. This was validated using an exhaustion flow cytometry panel at 40 weeks post-transduction.

**Results:** Using single-cell RNA-sequencing, CD8 pre-exhausted T-cells with self-renewing capacity (*TCF7*+), and terminally exhausted CD8 T-cells (*TCF7*-) were detected in the AAV-HBV model. These terminally exhausted CD8 T-cells (expressing *Pdcd1*, *Tox*, *Lag3*, *Tigit*) were significantly enriched versus control mice and independently identified through flow cytometry. Importantly, comparison to CHB human data showed a similar exhausted CD8 T-cell population in IT and IA donors, but not in healthy individuals.

**Conclusions:** Long term high titer AAV-HBV mouse liver transduction led to T-cell exhaustion, as evidenced by expression of classical immune checkpoint markers at mRNA and protein levels. In both IT and IA donors, a similar CD8 exhausted T-cell population was identified, with increased frequency observed in IA donors. These data support the use of the AAV-HBV mouse model to study T-cell exhaustion in HBV infection and the effect of immune-based therapeutic interventions.

**Lay Summary:** The AAV-HBV mouse model is used as a research tool to study hepatitis B infection. In this study we evaluated the translation value from mouse to human with regards to T-cell exhaustion.

**Highlights:**

- AAV-HBV mice transduced with a high titer vector showed presence of CD8 exhausted T-cells after 24 weeks.
- High titer transduced mice, but not lower titer show increased expression of LAG-3, TOX, TIM-3 and TIGIT in CD8 T-cells. PD-1 was increased in CD8 T-cells, independent of HBV transduction titer.
- A similar exhausted CD8 T-cell population could be found in chronic HBV donors, but not in healthy individuals.

**Graphical abstract:** 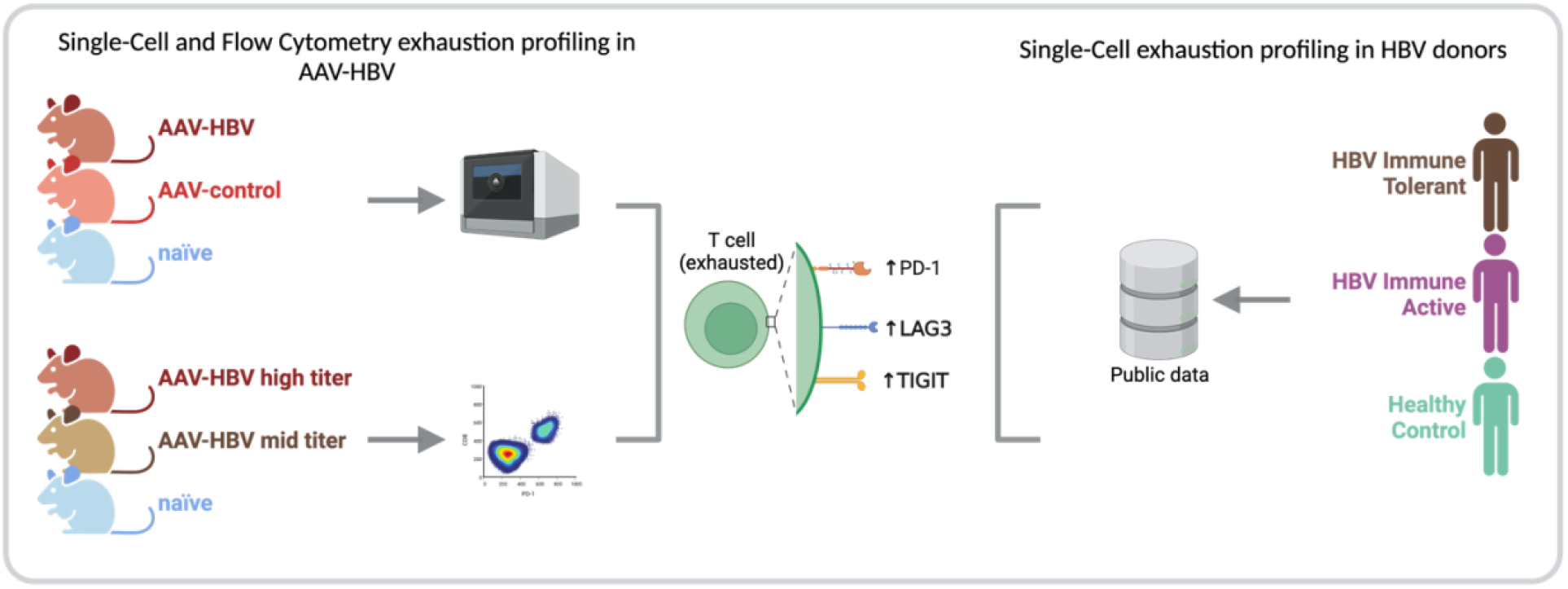

## Introduction

Hepatitis B virus (HBV) infection affects an estimated 296 million people worldwide, according to the World Health Organization (WHO) [1]. HBV is a double-stranded DNA virus from the *Hepadnaviridae* family [2] and primarily targets hepatocytes, leading to chronic liver inflammation and eventually, liver cirrhosis or hepatocellular carcinoma [3]. Despite the availability of prophylactic vaccines, vertical transmission still occurs, which leads to chronic infection in >90% of cases [1]. For those chronically infected, antiviral therapies control viral replication, but do not eradicate the virus even after many years. Functional cure is declared with at least 24 weeks of stable loss of detectable HBV DNA and surface antigen (HBsAg) after treatment cessation, since that duration predicts low relapse rates [2]. However, only a small fraction of patients achieve functional cure after treatment stop, likely due to lack of adequate immune control [4–6]. Even though the exact mechanisms underlying the impaired immune response aren’t fully understood, evidence of loss of CD4 T-cell help and dysfunctional HBV-specific CD8 T-cell responses have been reported [7, 8].

During acute viral infections, T-cell responses are believed to follow a pattern of expansion, contraction, memory cell formation and maintenance. However, in chronic infection, the continuous antigen exposure prevents proliferation-competent T-cells from returning to a memory quiescent state, which leads to an exhausted T-cell phenotype [9–12].

Exhausted T-cells were first described in lymphocytic choriomeningitis virus (LCMV) infection, which has a well-established mouse model of chronic infection. These exhausted T-cells were characterized by upregulation of several inhibitory receptors, such as PD-1, LAG-3, TIM-3, reduced cytokine production and diminished cytotoxic activity [13]. In chronic HBV, several studies have shown the presence of exhausted and dysfunctional T-cells [14–20]. The HBV-specific CD8 T-cells from chronically infected HBV patients, just like in cancer and other chronic viral infections, are not a functionally homogeneous population of exhausted T-cells, but distinct T-cell subsets with different degrees of dysfunction [21, 22] and present different HBV antigen specificities [23].

The AAV-HBV mouse model has emerged as an important tool to study HBV pathogenesis and as a preclinical model for drug discovery [24–26]. AAV-HBV transduction can lead to a long-term expression of viral antigens; however, the translational part of this model is still not well understood. We aimed to understand whether long-term AAV-HBV transduction in mice would lead to T-cell exhaustion. Moreover, we have analyzed publicly available data from human chronic HBV donors (immune tolerant (IT) and immune active (IA) phases) [27] to assess the similarity between the observed HBV T-cell exhaustion generated in the AAV-HBV mouse model and human CHB infection.

## Methods

### Ethical Statement and Animal Experimentation

Animal experiments were first reviewed, approved, and then conducted in strict accordance with guidelines established by the Janssen Pharmaceutica N.V. Institutional Animal Care and Use Committee. The local Johnson and Johnson Ethical Committee approved all experimental protocols performed at Janssen. Experiments were performed following the guidelines of the European Community Council directive of November 24, 1986 (Declaration of Helsinki 86/609/EEC). The criteria of the American Chemical Society ethical guidelines for the publication of research [28] were met. Every effort was made to minimize animal discomfort and to limit the number of animals used. Mice were kept in a specific pathogen-free facility under appropriate biosafety level following institutional guidelines.

### Study overview for single-cell RNA-seq

Male C57BL/6 mice (4–5-week-old, Wuxi Apptec, Shanghai, China) were either transduced via the tail vein with 1x10^11^ viral genome equivalents (vge) per mouse of rAAV-HBV1.3 (FivePlus Molecular Medicine Institute, Beijing, China) or rAAV vector control, which contained an empty rAAV capsid and rAAV containing a short sequence of the CMV promoter and BGH poly A tail flanked with inverted AAV-ITR. Twenty-four weeks post-transduction, samples from AAV-HBV mice (n=5) were collected based on persistent viral replication, with HBsAg titers >4 logIU/ml, along with mice transduced with AAV control vector (n=5) and naïve mice (n=5); control vector and naïve mice were randomly selected to provide an equivalent body weight distribution with the AAV-HBV mice.

### Study overview for flow cytometry

Female C57BL/6 mice (6-8 weeks old, Janvier labs, Le Genest-Saint-Isle, France) were transduced with rAAV8-1.3HBV (BrainVTA, Bejing, China) at 3x10^10^ vge/mouse (high dose) or 2.5x10^9^ vge/mouse (mid dose) via the tail vein. Twenty days after transduction mice were randomized based on HBsAg levels. Blood for viral parameters was collected via the saphenous vein every week, from which serum was prepared and stored at -80°C until assayed. Nine months post-transduction mouse liver samples were collected.

### Viral parameters

Serum HBsAg and HBeAg were quantified using CLIA kits (Cat. No. CL18002, CL18005, Ig Technology, Burlingame, CA, USA) which were used according to the manufacturer’s guidance. Dependent on the estimated levels, different dilutions of serum in PBS were used. Read-out of plates was performed with a Viewlux ultra HTS microplate imager (Perkin Elmer, Mechelen, Belgium).

### Mouse liver intrahepatic immune cell (IHICs) isolation for single-cell RNA-seq

IHICs were obtained by perfusing the liver with PBS via the hepatic portal vein to prevent contamination from circulating blood; single-cell suspensions of mouse IHICs were acquired via a mechanical dissection method using the GentleMacs (Milteny Biotec, Bergisch Gladbach, Germany). Homogenized liver tissue was centrifuged at 50 x *g* for 1 minute to decrease hepatocyte contamination. Cell suspensions were further purified with 35% Percoll (GE, 17-0891-09) in PBS and density gradient centrifugation at 800 x g for 20 minutes, followed by RBC lysis (eBioscience, 004333-57). Cell concentration and viability were then determined using a Countess® II Automated Cell Counter (Invitrogen).

### Mouse liver intrahepatic immune cell isolation for flow cytometry

IHICs were obtained by perfusing the liver with PBS via the hepatic portal vein to prevent contamination from circulating blood; tissue disruption was performed using a GentleMACS dissociator and the enzymatic liver dissociation kit for mice, according to the manufacturers protocol. Hepatocytes were separated from lymphocytes by centrifugation at 50× *g* for 5 minutes. Supernatant was spun down at 400× *g* for 5 minutes followed by resuspension in a an isotonic 33.75% (*v*/*v*) Percoll (GE Healthcare) diluted in PBS with 2% foetal calf serum and density gradient centrifugation at 700x *g* for 12 minutes. Next, residual hepatocytes and debris were discarded, and red blood cells co-sedimented with the IHICs were lysed using ACK lysis buffer (Lonza) for 5 minutes. Cells were washed twice and counted. Cell concentration and viability were then determined using a Nexcelom Cellaca MX Cell Counter (PerkinElmer). One million cells were used to perform flow cytometry staining.

To retrieve Kupffer cells and Liver sinusoidal endothelial cells (LSECs), an enzymatic digestion was performed for this study, instead of a mechanical digestion as performed for the single-cell RNA-sequencing study.

### Single-cell RNA-seq and data analysis

IHICs were processed according to the 10x Chromium Single-cell V(D)J Reagent Kits User Guide and loaded on to the Chromium 10x platform using the 5’ v1.1 chemistry (10x Genomics). Libraries were sequenced on a NovaSeq 6000 platform (PE150) (Illumina) to an average ∼50,000 reads per cell. Read alignment was done using the Cell Ranger pipeline (version 7) against the mouse genome reference (mm10). Resultant cell by gene matrices for each sample were merged across all conditions tested and samples. Pre-processing, alignment, and data filtering were applied equivalently to all samples using our internal pipelines based on OpenPipeline. Cells with less than 1,000 Unique Molecular Identifiers (UMIs) or less than 200 genes or more than 25% mitochondrial counts were removed from downstream analysis. All downstream analysis was done in R v4.0.5 [29] using the Seurat v4.1.1 package [30].

Data was log-normalized with a scaling factor of 10,000. The top 2,000 most variable genes as determined by the ‘vst’ method implemented as the FindVariableFeatures function were selected and scaled using a linear model implemented as the ScaleData function. Afterwards, principal component analysis (PCA) was run, the number of significant principal components (PCs) to be used for downstream cell clustering determined using an ElbowPlot and heatmap inspection. A nearest neighbor graph and Uniform Manifold Approximation and Projection (UMAP) plot were generated using the significant PCs.

A Louvain clustering was run on all cells, and the best resolution for clustering was determined using an average silhouette scoring across all clusters, testing 10 resolutions between 0.1 and 1 as previously implemented in Ziegler *et al* [31]. Marker genes for each cluster were calculated using the FindAllMarkers() function (method= ‘wilcox’) and each cluster was iteratively subclustered further using the same approach. Sub-clustering was stopped when the resulting clusters were not meaningfully different and no significant marker genes could be identified. Clusters were annotated as cell type populations based on the expression of their respective marker genes.

### Single-cell RNA-seq data differential expression analysis

A subpopulation pseudo-bulk analysis was performed using muscat v1.4.0 [32]. Each cell subpopulation raw data counts were aggregated to pseudo-bulk data using the aggregateData function with the fun=“sum” option. Differential state was assessed using the pbDS() function with the following parameters: method=“edgeR”, min_cells=1 (edgeR v3.32.1 [33]). Enrichment analysis was performed using the Gene Set Enrichment analysis of Gene Ontology (gseGO) function in the clusterProfiler v4.3 R package [34] (with an fdr adjusted P value <0.05).

### Single-cell RNA-seq data receptor-ligand analysis

After cell annotation, CellChat v1.1.3 [35] implementation in R v4.0.5 [29] was used to construct a cell– cell-communication network per condition. For naïve, AAV-control and AAV-HBV samples a network based on known ligand-receptor pairs and ligands was calculated using the standard analysis, omitting population size to decrease bias against low abundant cell populations.

### Human CHB dataset

Raw fastq files from the human chronic HBV dataset of Zhang *et al* [27] were downloaded from Gene Expression Omnibus (GSE182159). Read alignment was done using Cell Ranger 7.0.0 [36] against the human genome reference (GRCh38). Aligned cell by gene matrices for each sample were merged. All downstream analysis was done as described above for the mouse samples.

### Flow cytometry

All staining procedures were performed for 30 minutes at 4°C and all washing steps were done at 400× *g* for 5 minutes at 4°C. Cells were treated with True stain monocyte blocker (Biolegend) and an anti-CD16/CD32 FC blocking reagent (clone 2.4G2, BD) for 10 minutes, after which a dead cell exclusion marker (fixable viability dye eFluor780, Invitrogen) was co-incubated for 30 minutes. After washing the cells with stain buffer, staining was performed on ice using the following panel of fluorochrome-conjugated antibodies diluted in stain buffer (0.5% BSA (Bovine Serum Albumin) supplemented with brilliant stain buffer plus (BD)): CD45-BUV395 (clone 30-F11), CD8a-BUV496 (Clone 53-6.7), CD4-BV786 (clone GK1.5), CD3-PerCP-Cy5.5 (Clone 17A2), CXCR3-BV510 (clone CXCR3-173), PD1-BV605 (clone J43), LAG3-BUV737 (clone C9B7W), TIM3-BB515 (clone 5D12), (BD) and TIGIT-PE (cloneA17200C), CD154-PE-Cy7 (clone MR1), CD155-BV711 (clone TX56) (all mAbs from BioLegend). After staining of cell surface proteins, cells were washed twice with stain buffer, fixed with a fixation reagent (Invitrogen) and permeabilized twice using the Foxp3 / Transcription Factor Staining buffer set (Invitrogen) before being stained on ice with a second panel of antibodies added to diluted permeabilization buffer (BD): TOX-A647 (cloneNAN448B, BD), Foxp3-PE-CF594 (Clone 3G3, BD), TCF1-A405 (clone #812145, RnD systems). Finally, cells were washed twice with permeabilization buffer and left in 200µl of Stain buffer BSA in the dark at 4°C until cytometry data was acquired on a BD LSRFortessa instrument. Data were analyzed using FlowJo (BD).

### Statistical analysis

Statistical comparisons were performed using either GraphPad Prism 9 or using the package rstatix v0.7.2 [37] in R v4.0.5. Differential abundance in single-cell RNA-sequencing was calculated on cell proportions using a Wilcoxon test, and p-value adjusted using Bonferroni correction across all contrasts and populations.

## Results

### Intrahepatic immune cell atlas from AAV-HBV vs control

To build a murine HBV-specific immune cell atlas and characterize the T-cell exhaustion after prolonged HBV replication, we first characterized the AAV-HBV mouse model (and then compared to the human liver atlas samples from IT and IA CHB patients [27]). Liver samples from mice transduced with AAV-HBV (n=5), with AAV-control (n=5) or naïve mice (n=5) were collected 24 weeks post-transduction. All samples across groups showed good recovery of cells (Fig. S1) capturing a total of 79,970 high quality mouse immune cells and covering all major immune cell populations (Ts, Bs, NKs, DCs, monocytes and neutrophils) and hepatocytes (Fig. 1A-B). Kupffer cells made up 0.1% of the total cells, hepatocytes 5% while LSECs were not detected. In contrast, T-cells (52%) and B-cells (22%) were the most frequent cell types detected (Table S1). All major cell populations were well defined with hallmark genes (Fig. 1C) such as *Cd3d* for T-cells, *Cd79a* for B-cells, *Fapb1* for hepatocytes, *Cst3* for dendritic cells, *Lyz2* for monocytes/macrophages, *Fcerg1* for NK cells, *Cd63* for basophils, *S100a8/9* for neutrophils and *Prtn3* for endothelial cells. After performing the first level cell annotation analysis, each cluster was sub-clustered to obtain greater granularity (Fig. S2). B-cells were further subdivided into plasma cells, pre B-cells and mature B-cells (Table S2). Within the T-cell compartment, a first subclustering identified *CD8*, *CD4* and *CD3+* positive T-cells and each T-cell subtype was further defined based on key marker genes (Fig. 1C, Fig. S3, Table S2). The dendritic cells were subdivided into pDCs, cDC1 and cDC2. The monocyte compartment contained classical monocytes, patrolling monocytes and proliferating monocytes; the macrophages identified were Kupffer cells and capsule macrophages. For the NK cells, a liver-resident NK population was identified. Finally, the neutrophils were subdivided into mature neutrophils, type I interferon neutrophils (Neutrophils-IFN) and immature neutrophils.

**Fig. 1.**
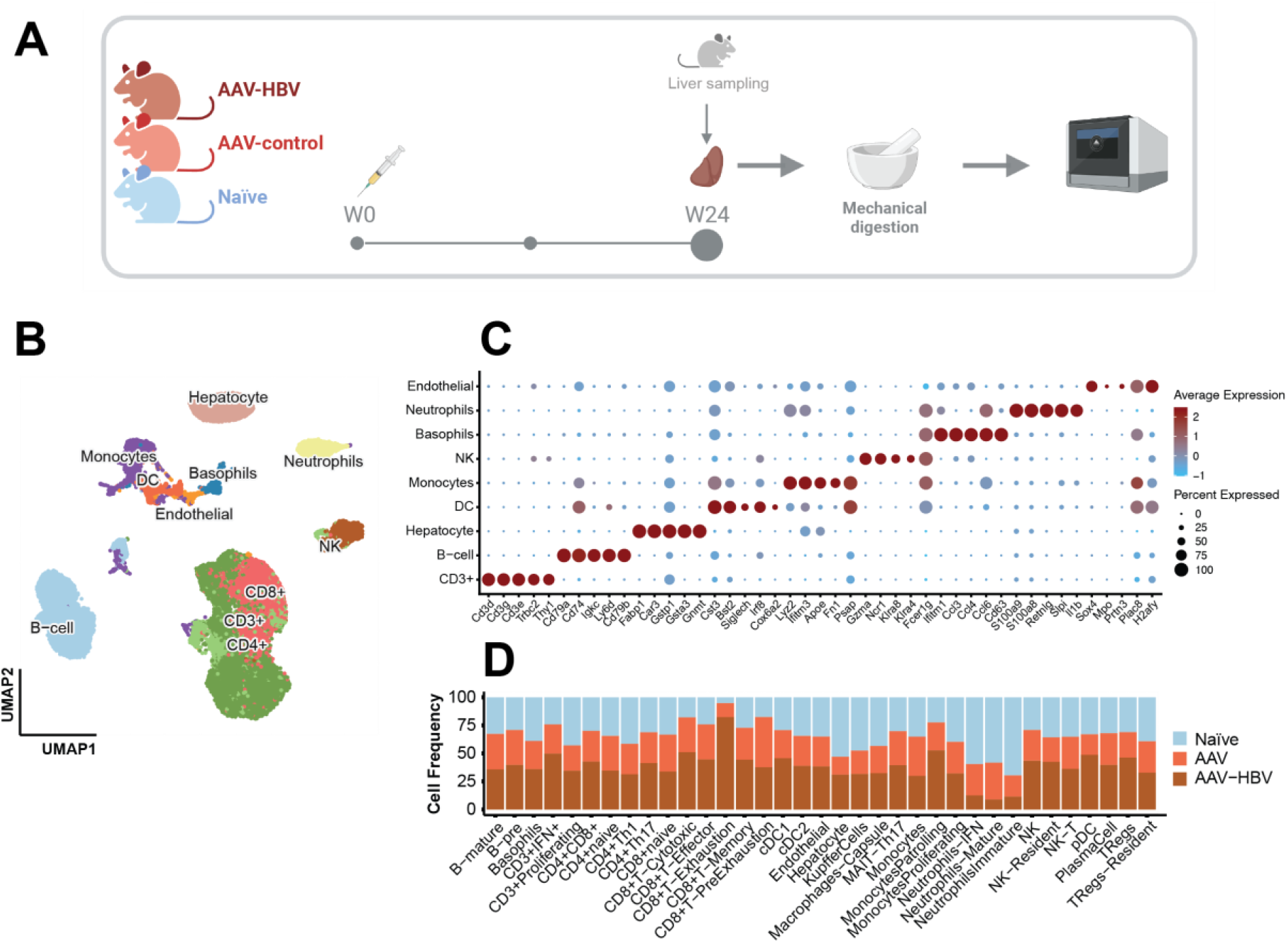
Single-cell RNA sequencing profiling of the AAV-HBV mouse model. Liver samples from male C57/bl6 mice from three groups were collected: infected with AAV-HBV, with AAV-control or naïve 24-weeks post transduction. **(A)** Graphical overview of the experimental design using liver mechanical digestion for intrahepatic lymphocyte isolation and single-cell RNA-sequencing profiling. **(B)** UMAP projection of the major cell types identified from 15 samples. **(C)** Dotplot of scaled data with expression of the top 5 marker genes across each major cell type, determined using Wilcoxon Rank sum test. **(D)** Cell proportion (%) plot across all cell types (deepest cell annotation) in the three different mouse groups.

All cell subpopulations could be identified across the three groups (AAV-HBV, AAV-control and naïve mice) (Fig 1D). No statistically significant cell proportions differences between AAV-HBV and AAV-control samples were observed at the highest level of cell annotation (Table S3). Deeper analysis of the T-cell compartment (CD8+, CD4+ and CD3+) between AAV-HBV and AAV-control samples showed no differences among CD4 and CD3 in presence/absence of HBV replication (Table S4). Interestingly, this was different for CD8 T-cells, where 75% of the CD8 exhaustion T-cells were observed in the AAV-HBV arm (Fig 1D, Table S4).

### Prolonged HBV replication induces an expansion of CD8 T-cells with an exhaustion phenotype in the AAV-HBV mouse model

After 24 weeks of AAV-HBV transduction and HBV replication, the CD8 T-cell compartment showed a broad range of cell types (Fig. 2A and 2B), from naïve to more effector states across HBV and control arms. CD8 naïve T-cells expressed high levels of *Tcf7*, *Sell* and *Lef1;* CD8 naïve to memory cells also expressed the same naïve makers, but at lower levels, and high levels of *Ly6c2* [38]. The cytotoxic CD8 T-cells expressed granzyme B (*Gzmb*), *Cd160* and *Xcl1* [39]. The effector CD8 T-cells expressed the well described *Cx3cr1* [40]. Interestingly, two sub-populations, labeled pre-exhaustion T-cells (Tpex) and exhausted T-cells (Tex) were identified.

**Fig. 2.**
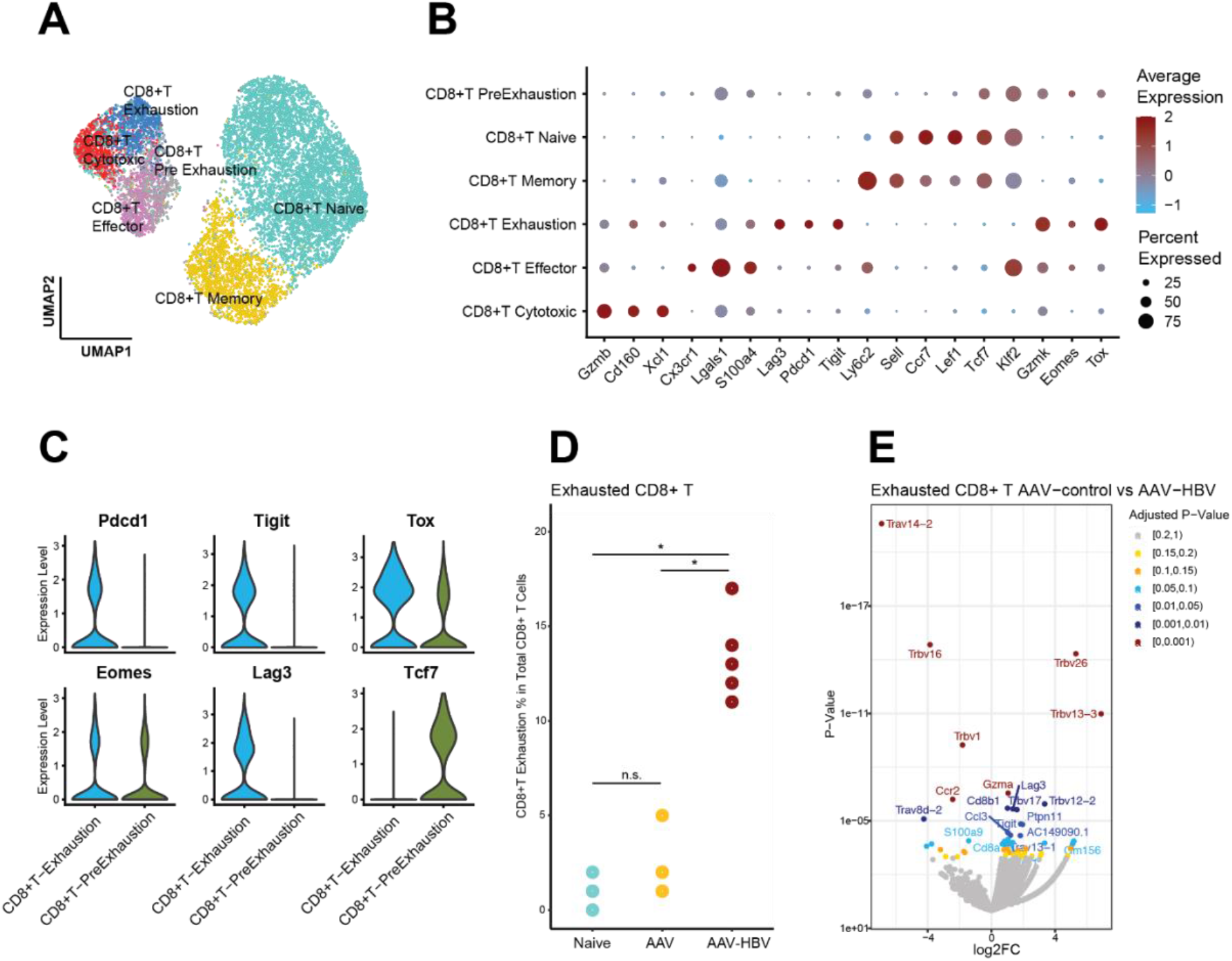
Single-cell sequencing demonstrates CD8 T-cell exhaustion in AAV-HBV mouse model. **(A)** UMAP projection of the deeper annotation of all CD8 T-cell populations across the different mouse groups. **(B)** Dotplot with expression of the top 3 marker genes across each deepest CD8 T-cell annotation. **(C)** Violin plots of log normalized data from the exhaustion markers *Pdcd1*, *Tigit*, *Tox*, *Eomes*, *Lag3* and *Tcf7* across the exhausted T-cells and pre-exhaustion T-cells. **(D)** Frequency of the CD8 Exhausted T-cells across the different mouse groups. Statistical comparisons were done using a non-parametric Wilcoxon test and Bonferroni correction. Comparisons with adjusted p-value <0.05 are shown with (*). **(E)** Volcano plot showing the differently expressed genes calculated using edgeR between AAV-control and AAV-HBV in CD8 Exhausted T-cells. Positive log_2_ fold-changes indicate higher expression in AAV-HBV. Genes are colored by adjusted p-value.

The Tpex population (Fig. 2C), showed expression of checkpoint inhibitor markers (*Tox* and *Eomes*) and *Tcf7* (encoding for TCF-1, which is lost upon transition to a fully exhausted phenotype [41]). On the other hand, the Tex cells showed expression of *Pdcd1* (encoding for PD-1), *Tigit*, *Tox*, *Eomes* and *Lag3*, but no *Tcf7* expression (Fig. 2C). Differential expression analysis within the Tex cells across AAV-control and AAV-HBV showed significantly increased *Tigit*, *Lag3* and *Gzma* expression in the AAV-HBV versus the AAV-control arm showing the observed exhausted phenotype was not related to the AAV component, but due to HBV transduction (Fig. 2E). Compositional analysis across the three groups showed that all AAV-HBV mice had over 10% of their total CD8 T-cells as Tex, with an average of 13.4% (SD ±2.3), whereas in the control arms (both AAV-control and naïve mice) most CD8 T-cells were phenotypically functional as Tex were only on average 2.2% (SD ±1.6) and 1.0% (SD ±0.7), respectively. When testing for differential abundance on the Tpex population, both AAV-control (Wilcoxon rank-sum test, p-value=0.036) and naïve (Wilcoxon rank-sum test, p-value=0.033) groups were significantly different from AAV-HBV, but not AAV-control vs naïve (Wilcoxon rank-sum test, p-value=0.525) (Fig 2D). The abundance of Tpex cells was not significantly different across any of the groups (Fig. S5).

### High HBV antigenic titer correlates with higher level of CD8-T cell phenotypic exhaustion

An independent study using flow cytometry as a read out was performed to validate the single-cell findings demonstrating T-cell exhaustion in the AAV-HBV mouse model 24 weeks post-transduction and to assess whether this phenotype was influenced by the level of HBV antigenemia. C57BL/6 mice (n=6 mice/group) were transduced with either a high titer of AAV-HBV (3x10^10^vge/mouse), a lower mid-titer of AAV-HBV (2.5x10^9^ vge/mouse) or kept naïve. At the start of the study the high titer mice had a mean HBsAg level of 4log_10_IU/ml, the low titer mice 3log_10_IU/ml, at the end of the study the HBsAg titers in serum were respectively 3log_10_ IU/ml and 2log_10_IU/ml (Fig. S4). To evaluate our goal, IHIC were stained for relevant markers identified by single-cell RNA-seq: PD-1, LAG-3, TIGIT, TOX and TCF-1. This gene set was supplemented with TIM-3, a well-known checkpoint inhibitor marker, even though the gene (*Havcr2*) was not detected in the AAV-HBV exhausted population.

An initial analysis was performed to compare the proportions Tpex and Tex in both flow cytometry and single-cell RNA-sequencing. Flow cytometry from the high titer mice group showed an average of 4.6% (±1.7 SD) of Tex in CD8 T-cells, characterized by PD-1+, LAG-3+ and TCF-1-(Fig. 3A) whereas Tpex within CD8 T-cells made up 1.6% on average (±1.9 SD), characterized by TOX high and TCF-1+ (Fig. 3B). The single-cell Tex (available from high titer mice) had an average abundance of 14%, whereas the Tpex had an average of 6.2% (±2.2 SD). The proportion of exhausted T-cells detected by flow cytometry was lower compared to those detected by single-cell RNA-sequencing (Fig. 3B). However, the relative proportions of the 2 populations (Tex and Tpex) were comparable between the two different techniques, where higher proportions of both Tex (3-fold higher) and Tpex (3.8-fold higher) were observed in the single-cell dataset (Table S5).

**Fig. 3.**
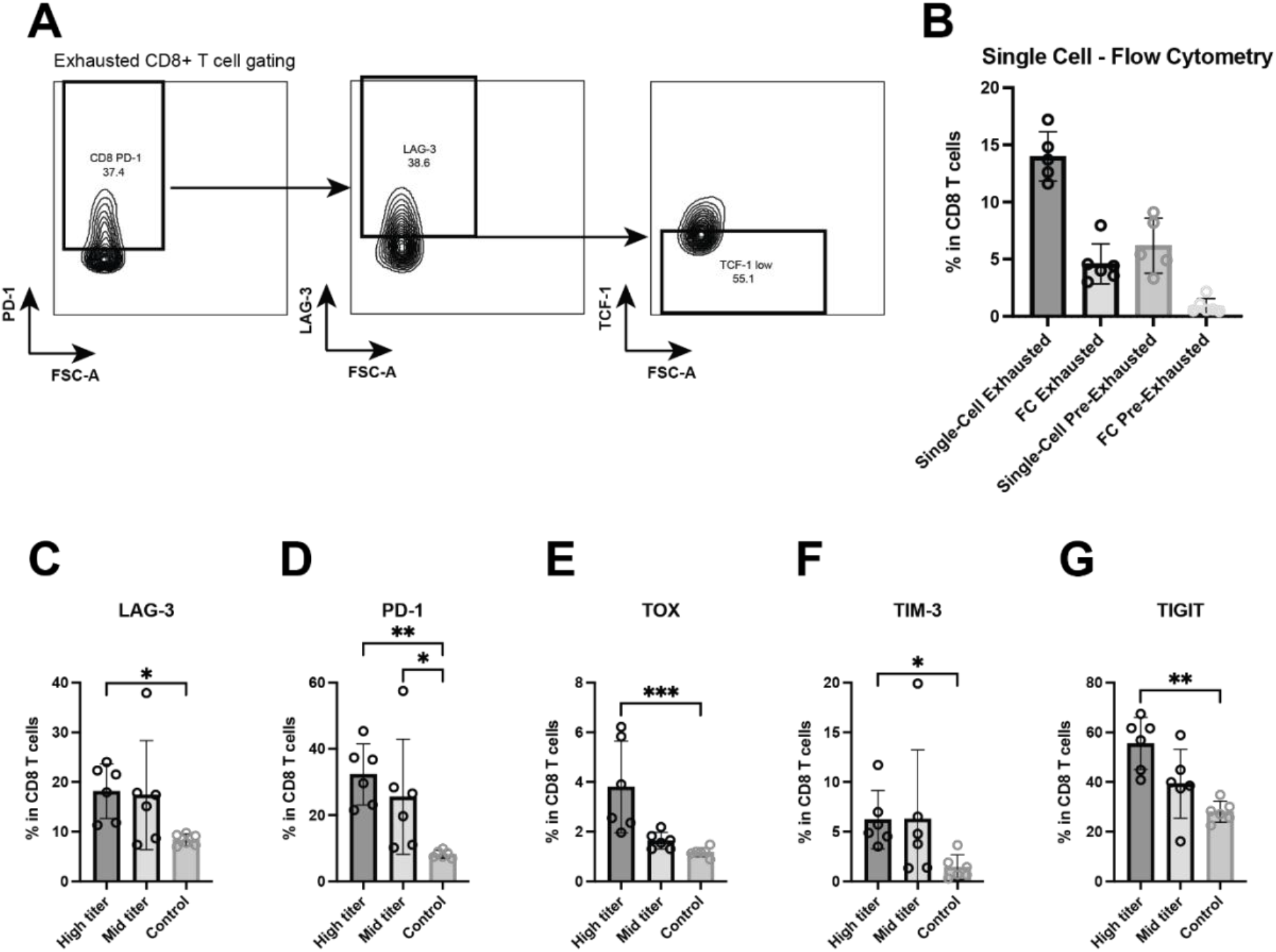
Flow cytometry demonstrates exhaustion marker expression on liver CD8 T-cells isolated from AAV-HBV transduced mice. **(A)** Representative plot of CD8 T-cells that are PD-1+, LAG-3+ and TCF-. **(B)** Percentage of exhausted and pre-exhausted T-cells within the CD8 T-cells detected by single-cell RNA-sequencing or by Flow Cytometry. **(C)** LAG-3+, **(D)** PD-1+, **(E)** TOX+, **(F)** TIM-3+ and **(G)** TIGIT+ CD8 T-cells in the AAV-HBV mice dosed with high titer of virus or mid titer of virus. Data are shown as mean with errors (SD) from a single experiment run with group sizes of 7. A Kruskal-Wallis test was performed across the three groups and significant comparisons are highlighted.

The evaluation of expression of the different exhaustion markers showed clear differences depending on the HBV antigenic titer (Fig. 3C-G). The percentage of T-cells in the CD8 compartment positive was significantly increased in high titer mice vs naïve (but not in mid-titer mice vs naïve) using a Kruskal-Wallis test for LAG-3 (p=0.013), PD-1 (p=0.002), TIM-3 (p= 0.018), TIGIT (p= 0.003) and TOX high (p= 0.001), with TOX showing the highest significance (Fig. 3C-G). Only the expression of PD-1 was significantly higher between the mid titer mice compared to the naïve controls (Kruskal-Wallis test, p=0.039); the other markers were not significantly different between these two groups. To conclude, Tex and Tpex populations were detected on the protein level and expression of most exhaustion markers increased in association with a higher serum HBsAg titer suggesting that higher HBV antigenemia leads to a higher level of T-cell exhaustion.

In the CD4 T-cell compartment the frequency of both regulatory T-cells (Treg) and Tfh cells was significantly greater in the high titer group compared to the control mice (Kruskal-Wallis test, p=0.001 for Tregs and p=0.033 for follicular helper, Fig 4A). CD4 T-cells did express exhaustion markers. However, there was no difference in expression between high titer and mid titer mice (Fig. 4B-E). PD-1 and TIGIT positive CD4 T-cells both showed significantly greater frequencies between high titer mice and controls (Kruskal-Wallis test, p= 0.01 and p=0.002, respectively, Fig 4C-D). The CD4 Tfh were not detected in the single-cell RNA-sequencing data, whereas with flow cytometry they made up on average <1.5% of the total CD4 population for all groups (Fig. 4A).

**Fig. 4.**
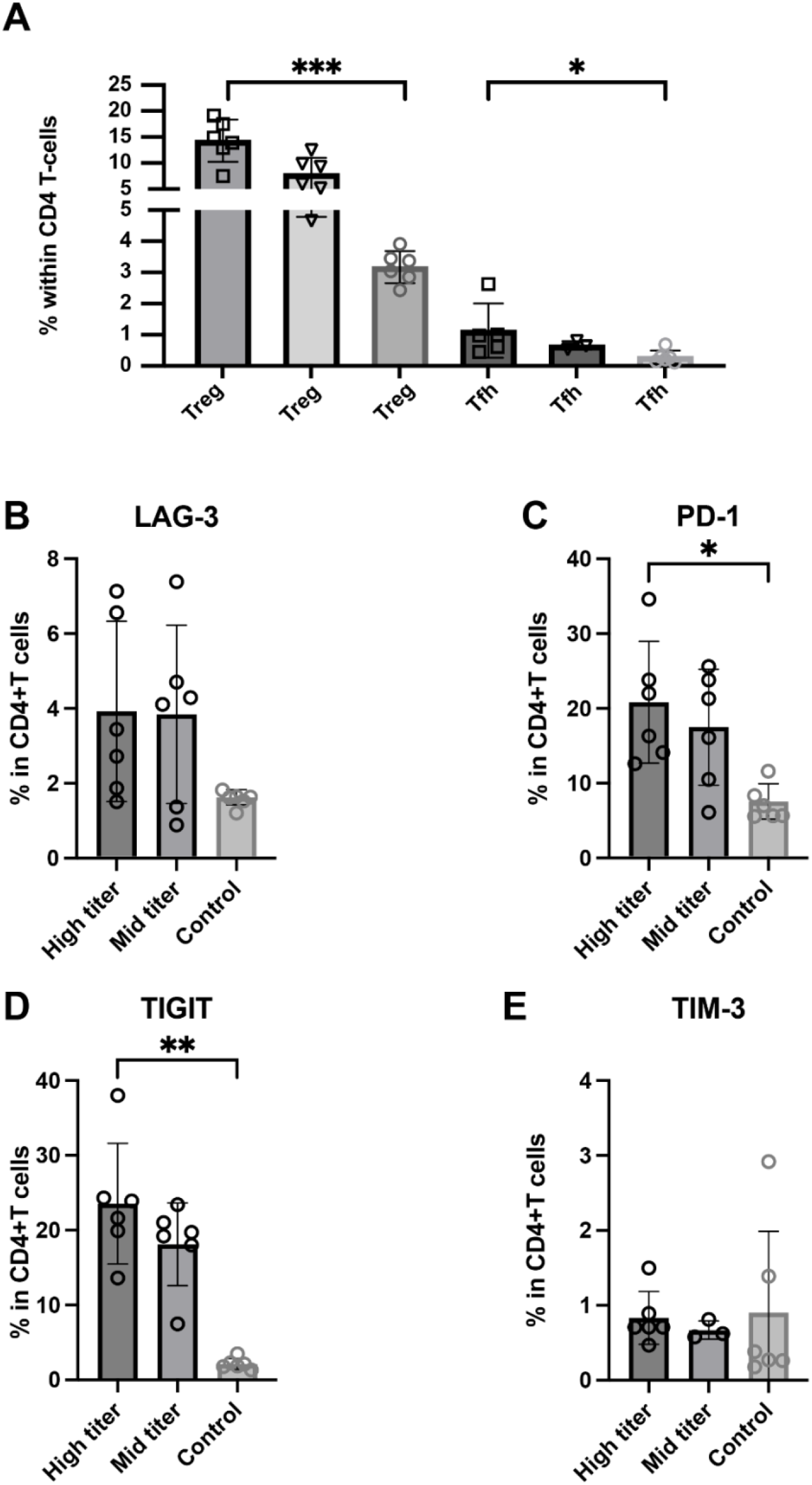
Flow cytometry demonstrates expression of exhaustion markers on liver CD4 T-cells isolated from AAV-HBV transduced mice. **(A)** Percentage of regulatory T-cells and follicular helper T-cells within the CD4 T-cell population. **(B)** Percentage LAG-3+, **(C)** PD-1+, **(D)** TIGIT+ and **(E)** TIM-3+ CD4 T-cells. Data are shown as mean with errors (SD) from a single experiment run with group sizes of 7. Kruskal-Wallis test was performed across the three groups and significant comparisons are highlighted.

### Human atlas and analyses shows translatability of AAV-HBV model CD8 T-cell exhaustion with IT and IA CHB stages of disease

Using the same cell typing approach a human liver atlas was built and similar analyses identified a CD8 T-cell exhausted population in human liver samples from IT and IA CHB donors [27]. The human Tex cells expressed *PDCD1*, *TOX*, *TIGIT*, *GZMK*, *LAG3* and no *TCF7*, similarly to the mouse Tex (Fig. S6). Additionally, the human Tex cells also expressed *LAYN* and *CTLA4*, which were not detected in mouse cells (Fig. S6).

After identification of cell types, using an iterative subclustering approach, the frequencies of Tex in human liver across the different HBV disease stages were assessed and compared. The IA donors showed increased frequency of Tex compared to IT (Wilcoxon rank-sum test, p=0.033) donors and healthy control donors (HC) (Wilcoxon rank-sum test, p=0.019). This increased frequency of Tex was also observed in the AAV-HBV model after prolonged (24 week) transduction time (Fig. 5A).

**Fig. 5.**
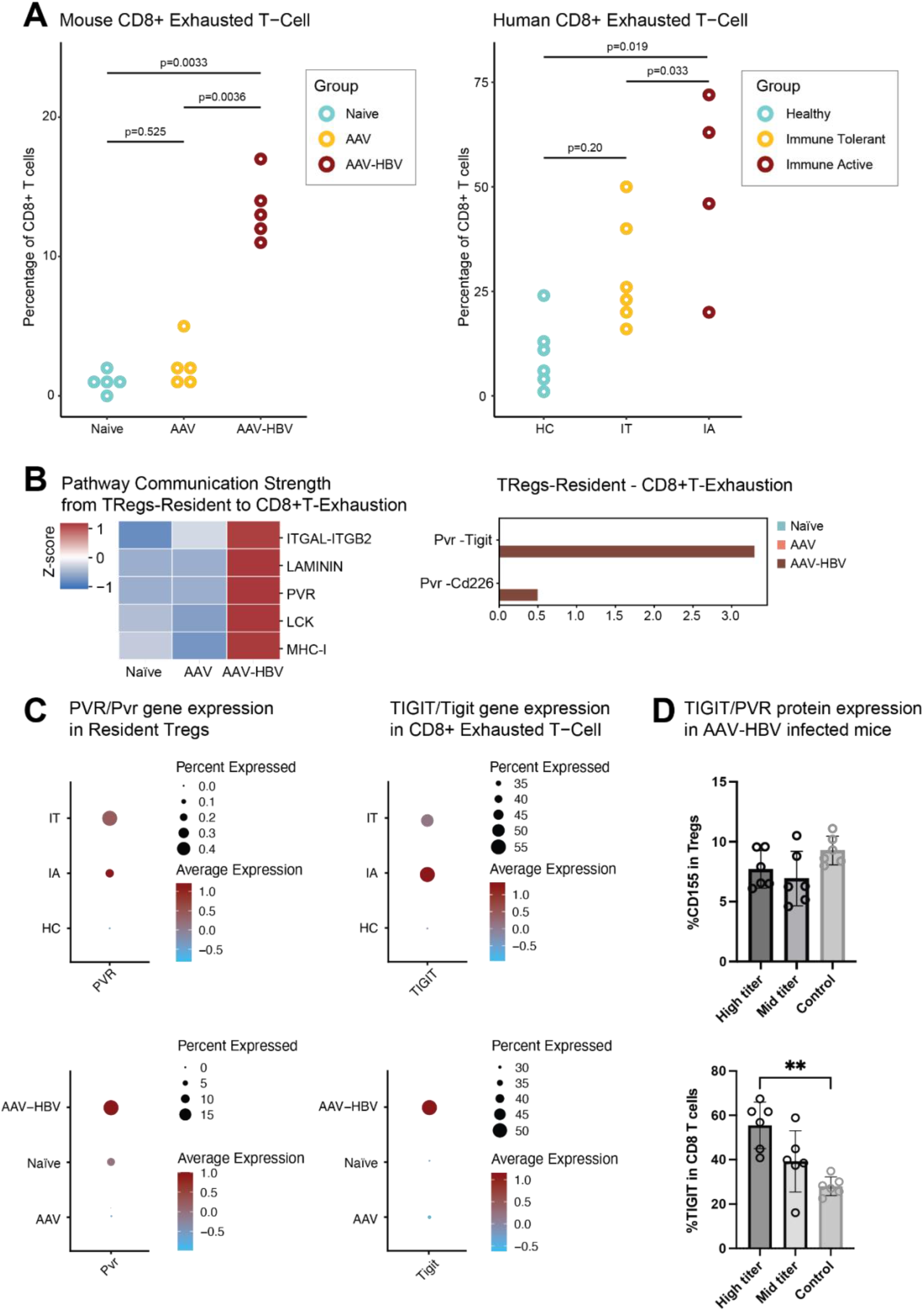
CD8 exhaustion is present at mRNA and protein level in the liver of AAV-HBV transduced mice and at mRNA level in human HBV liver samples. (A) Proportion of the CD8 Exhausted T-cells across the mouse and human. The human samples are divided in healthy controls (HC), immune tolerant (IT) and immune active (IA) donors. Statistical comparisons were done using a non-parametric Wilcoxon test and Bonferroni correction. (B) Heatmap from the cell-cell communication strength z-score for pathway communication between resident Tregs and exhausted CD8 T-cells. Barplot for the *Pvr* pathway interaction strength between the different mouse groups. (C) Dotplot of scaled data of the gene expression of *Pvr*/*PVR* and *Tigit*/*TIGIT* across the different mouse and human sample groups. (D) Barplot with the frequency of protein expression from flow cytometry analysis. Percentage of CD155 (PVR) in the Tregs and percentage of TIGIT protein expression in CD8 T-cells from AAV-HBV infected mice. Data are shown as mean with errors (SD) from a single experiment run with group sizes of 7. Statistical comparisons were done using a non-parametric Mann Whitney test. Comparisons with p-value <0.05 are shown with (*).

To understand the potential of cell-cell interactions between Tex and other cell types, we used CellChat, which makes use of a pre-defined database of receptors and ligands. The algorithm predicts significant communications, by identifying differentially over-expressed ligand and receptor pairs for each cell population [35]. Deconvolution of cell-to-cell signaling pathways showed that resident Tregs showed strong communication strength with the CD8 Tex cells (Fig. 5B). After identification of this cell-cell interaction, the pathways driving the cell-cell communication were explored (Fig. 5B), identifying PVR (poliovirus receptor) as an up-regulated pathway. This pathway encompassed both Pvr-Tigit and Pvr-Cd226 receptor-ligands (Fig. 5B). Interestingly, the Pvr-Tigit interaction between resident Tregs and CD8 Tex, showed an increased communication score in AAV-HBV, but not in AAV control nor naïve mice (Fig. 5B).

The follow-up study in AAV-HBV mice using flow cytometry showed that PVR detection in Tregs from mice infected with high (7.7% ±1.6 SD), mid-titer (6.9% ±2.3 SD) and controls (9.3% ±1.2 SD) was observed (Fig. 5D). No significant differences were observed across the three groups. However, within the CD8 compartment, a significant increase in the proportion of CD8s expressing TIGIT was observed in the high titer compared to the control naïve mice (Fig. 5D).

In CD8 Tex cells of the AAV-HBV arm, an increased average expression of *Tigit*, together with increased *Pvr* expression in resident Tregs (Fig. 5C) was observed. Across the human samples, *TIGIT* was only expressed in the IT and IA samples, but not in healthy controls nor in chronic resolvers (Fig. 5C). The same was observed in the Tregs and *PVR*, where *PVR* expression was only seen in IT and IA samples.

## Discussion

In this study a mouse and human liver atlas of intrahepatic T-cells was constructed using single-cell RNA-sequencing data from a long (24 week) AAV-HBV transduction model as well as from samples from CHB at IT and IA stages of infection. The single-cell observations were validated by flow cytometry on AAV-HBV mouse IHICs and the correlation of the findings to human data was confirmed. Our approach to isolate IHICs from the liver using mechanical digestion resulted in good recovery of the T-cell and B compartments but impacted the recovery of fragile CD45-fraction of cells, such as Kupffer cells and LSECs. Their recovery could likely be improved by using methods implementing an enzymatic digestion [42]. Though not significantly different, the uneven distribution of neutrophils across study groups might be due to technical capture issues of the technology (Fig. 1D, Table S3) as previously described [39].

Chronic viral infections lead to T-cell exhaustion and the LCMV mouse model has been extensively used to investigate this phenomenon. In contrast, the extensively used AAV-HBV mouse model, lacks an in-depth characterization of the immune liver environment and, in particular, the T-cell exhaustion levels and phenotype of the AAV-HBV mouse model are still unknown. Studies in LCMV have identified heterogeneity in Tex phenotypes which are characterized by distinct surface receptors, functionality, proliferative capacity, and tissue localization during chronic viral infections [13, 43–45]. Interestingly, the pool of Tex has been shown to be replenished by precursor-like T-cells (Tpex) that exhibit self-renewal capacity and are TCF-1 dependent. Tpex are characterized as PD-1+, TOX+ and TCF-1+ (TCF-7 gene), whereas Tex are characterized as PD-1+, TOX+ and TCF-1-[44]. Similar to the LCMV model, we found that Tpex and Tex in liver samples from AAV-HBV transduced mice, though the Tpex pool was similar in the AAV-HBV and AAV control arms. Tpex cells have been described as critical to maintain the pool of exhausted T-cells [46]. The transcription factor TOX might be critical to identify this Tpex population, since absence of TOX has been described to lead to a reduction in the Tpex population [47–49]. In this study an increase in TOX expressing CD8 T-cells was observed in mice transduced with a higher AAV-HBV titer, suggesting that these might be responsible for the establishment and maintenance of the exhaustion phenotype observed in the (high titer) AAV-HBV model. Moreover, in the absence of antigen it is postulated that Tpex cells transition into conventional memory-like cells with self-renewing capacity [44, 46]. Importantly, we used *Tcf7* (encoding TCF-1) to distinguish between terminally exhausted T-cells and the pre-exhausted phenotype (Fig. 2C). TCF-1 has been linked to cell renewal capacity and effector function in chronic infection, which is lost upon transition to a fully exhausted phenotype [41, 46, 50, 51]. To determine the molecular signature of exhaustion, frequency profiles of IHICs from control and AAV-HBV mice were compared. These data showed that within CD8 T-cells, Tex, but not Tpex could be identified in higher frequency in AAV-HBV infected mice compared to control mice, suggesting that by 24 weeks post-transduction, CD8 T-cells have become phenotypically exhausted in the AAV mouse model.

The exhaustion phenotype is a continuous phenotype [52, 53], and it is challenging to categorize different states of exhaustion with flow cytometry, as we are limited to a smaller set of protein markers. This is a limitation of the technology, even though it allows for a simple cell type delineation based on protein marker association. It is worth pointing out that, in the follow up study, flow cytometry showed the presence of Tex, in the high titer AAV-HBV arm though at a lower frequency than through single-cell RNA-sequencing (it is well established that mRNA expression only moderately correlates with protein expression [54]). A study has shown that in CD8 T-cells, the differences between mRNA and protein are independent of T-cell differentiation or activation status [54]. Moreover, the assignment of Tex in flow cytometry gating is done on a cell-by-cell level, whereas single-cell RNA-seq is done based on clustering approaches and could inflate frequencies. Nevertheless, and importantly, the ratio between Tex and Tpex was maintained regardless of the technology, with a greater proportion ratio of Tex/Tpex being detected (Fig. 3B). The proportion of cells expressing PD-1 at protein level was significantly increased between both high and mid-titer versus controls. For the remainder of the markers, only the high titer was increased versus controls, with TOX showing the highest increase and significance (Fig. 3C-G). Apparently, a higher titer AAV-HBV transduction induces a more pronounced exhaustion phenotype. Moreover, since the exhaustion phenotype could be observed both in single-cell RNA-sequencing at 24 weeks and in flow-cytometry at 9 months (40 weeks) with the high titer transduction, but not with the mid titer transduction, it suggests that titer rather than time post-transduction (i.e HBV replication) is key. Interestingly, it has been described in a study looking at HBV-specific CD8 T-cells from HBV donors, that TOX levels were higher in chronic HBV donors versus resolved patients [55]. Moreover, in LCMV, it has been described that TOX expression is dependent on the antigen dose, and it promotes T-cell dysfunction and long-term survival [48]. Finally, our data showed a significantly higher proportion of CD4 Tregs in high titer mice compared to lower titer mice (Fig 4A). This is in concordance with human data were higher frequency of Tregs was observed in chronic HBV-infected individuals versus healthy controls [56]. Furthermore, inhibitory receptor molecule expression on liver-resident CD4 T-cells from AAV-HBV transduced mice is comparable to those described in literature pertaining to humans / patients [57].

To the best of our knowledge, this is the first time that single-cell RNA-sequencing has been performed on IHICs from AAV-HBV infected liver samples and used to compare with human HBV samples. A previous study in a different mouse model (using Alb-Cre transgenic mice transduced with AAV-rcccDNA) demonstrated the presence of exhaustion after 8 weeks [58]. This exhausted population also expressed *Pdcd1* and *Tox*, but only reached a frequency of 1.49% of total CD8 T-cells [58]. The difference between that published study and our data may be due to either the shorter period of transduction, or a milder exhaustion phenotype in this model.

It has been described that peripheral T-cells from HBV-infected individuals are exhausted [14, 16, 59]. Moreover, a recent study has investigated the immune landscape in liver of chronic hepatitis B virus patients from different stages of disease at the single-cell level [27]. These liver biopsy data were re-analyzed, and the same exhausted population could be detected in IT and IA HBV-infected individuals, showing that the immunological phenotype of the AAV-HBV mouse model is analogous to the human disease. This provides confidence in the use of this model for the evaluation of therapies interfering with immune exhaustion pathways.

More recently, tools to further understand intercellular communications from single-cell RNA-sequencing data have been developed [35, 60]. Zhang *et al* described a potential interaction between Tregs and Tex, and increased frequency of both population in IT and IA donors and we aimed to understand the same in the AAV-HBV model [27]. The analytical cell-cell communication tool CellChat, encompasses a manually curated database of ligand-receptor interactions, which were corralled into pathways. An interesting correlation between Tregs-Tex and the PVR-TIGIT axis was observed. It is well established that *TIGIT* is expressed in T-cells, such as exhausted T-cells and Tregs. *PVR* was thought to be expressed in APCs and tumor cells [61]. In the analyzed data, even though *Pvr*/*PVR* was expressed in a small percentage of the Tregs from both AAV-HBV mice and IT and IA human samples, it was a consistent observation across the two studies (Fig 5C). We used flow cytometry in AAV-HBV mice, to further validate this finding, and observed that a comparable percentage (7-8% PVR expression in Tregs) of protein expression was observed [61]. The fact that a small percentage of Tregs expresses PVR might have implications towards cell-cell interaction and warrants further investigation. It remains to be seen whether these Tregs can further enhance the immunosuppressive phenotype of Tex. Finally, Tregs were significantly increased in frequency in high titer mice vs controls in the flow cytometry assessment but not in the single-cell analysis (possibly due to low cell numbers in single-cell), which was also described in IT and IA donors [27]. Importantly, the levels of antigenemia (i.e. HBsAg levels) in the high titer AAV-HBV mouse model is more representative of the levels observed in CHB patients, providing confidence about the translatability of the described liver immune environment.

Taken together, the results of this study confirm that the (high titer) AAV-HBV mouse model 6 months post-transduction induces a T-cell exhaustion phenotype similar to that seen in CHB patients. Interestingly, the plasma concentrations of HBsAg and HBeAg have a clear effect in the upregulation of exhaustion markers TIGIT and TOX protein expression levels, suggesting a higher exhaustion burden and less functional T-cells when those antigens are high. In chronic HBV patients it has been shown that very low HBsAg levels (<100 IU/ml) are associated with spontaneous HBsAg clearance due to functional and less exhausted T-cells in comparison to those with high HBsAg levels [62].

To conclude, by comparing liver samples from human CHB patients and AAV-HBV transduced mice we demonstrated that the CD8 T-cell exhaustion phenotypes were analogous. These data support the use of the AAV-HBV mouse model for the study of potential therapeutic approaches towards the reversal of T-cell exhaustion in chronic HBV infection.

## Acknowledgments

We would like to thank Frederik Pauwels for his support on the study and Heather Davis and Richard May for helpful comments on the manuscript.

## Abbreviation list

CHB: chronic hepatitis B
HBV: hepatitis B virus
HC: healthy controls
IA: immune active
IHIC: intrahepatic immune cell
IT: immune tolerant
LCMV: lymphocytic choriomeningitis virus
LSEC: Liver sinusoidal endothelial cells
PVR: poliovirus receptor
Tex: exhausted T-cells
Tpex: pre-exhaustion T-cells
UMAP: Uniform Manifold Approximation and Projection
UMI: Unique Molecular Identifier
vge: viral genome equivalents
WHO: World Health Organization

