## SupplementaryFiles for "AAV-HBV mouse model replicates immune exhaustion patterns of chronic HBV patients at single-cell level"

**SUPLEMMENTARY FIGURES**

**Fig. S1**

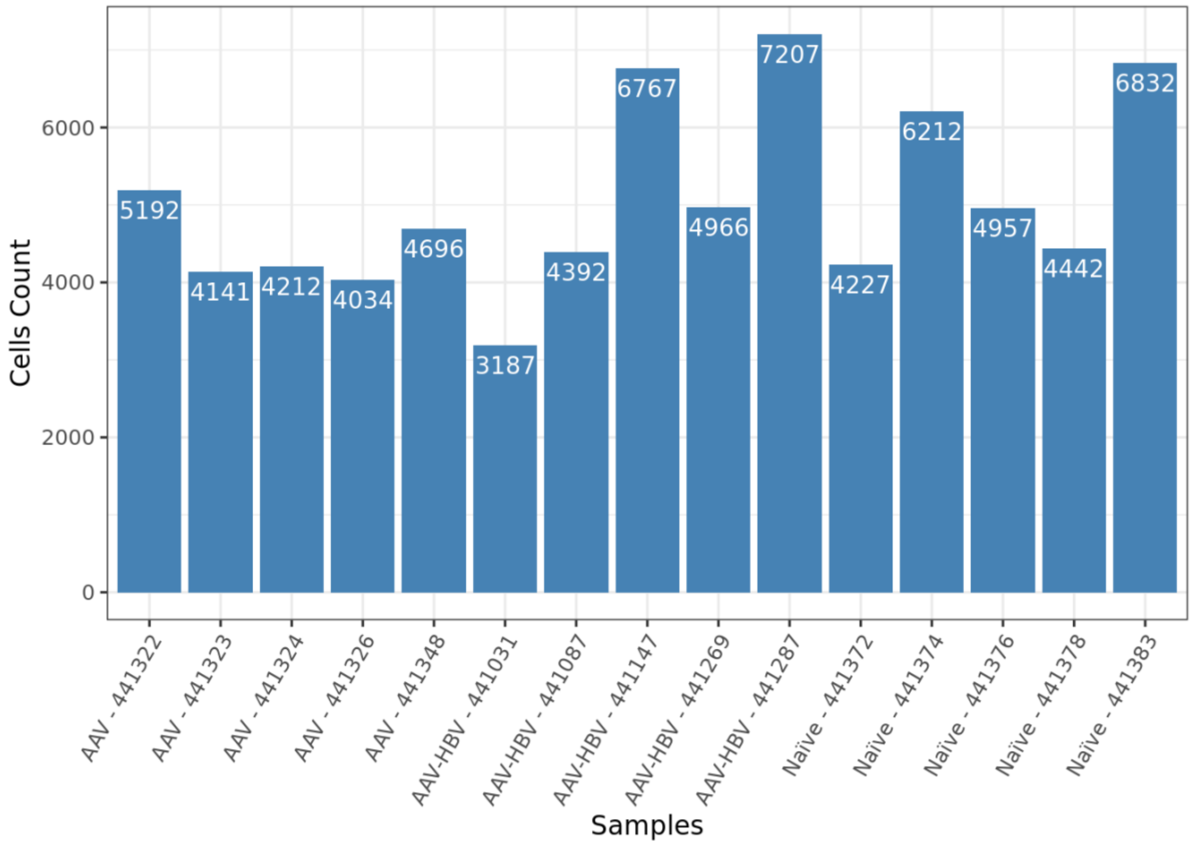

**Fig. S2**

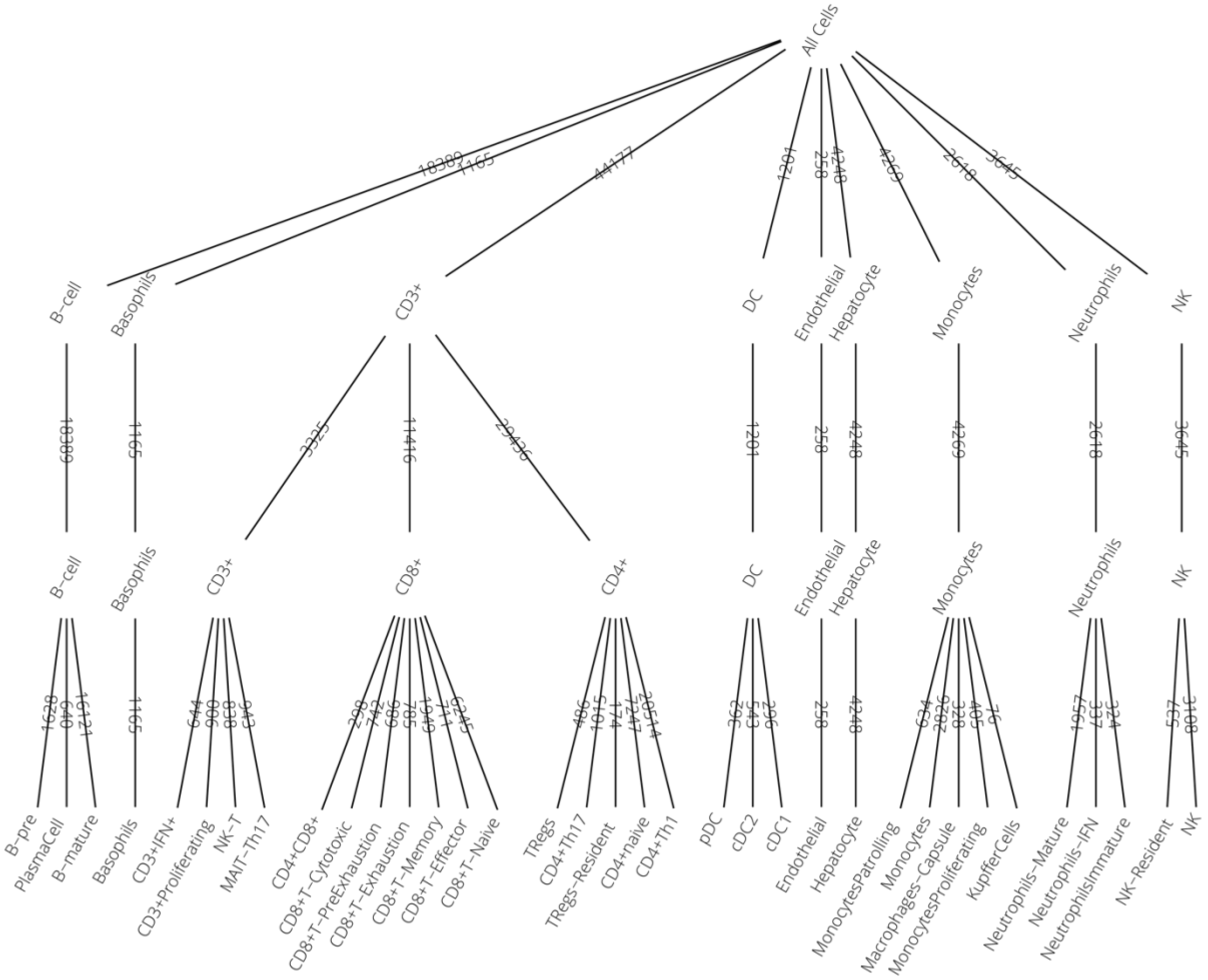
All Cells
B−cell
Basophils
CD3+
DC
Endothelial
Hepatocyte
Monocytes
Neutrophils
NK
B−cell
Basophils
CD3+
CD8+
CD4+
DC
Endothelial
Hepatocyte
Monocytes
Neutrophils
NK
B−pre
PlasmaCell
B−mature
Basophils
CD3+IFN+
CD3+Proliferating
NK−T
MAIT−Th17
CD4+CD8+
CD8+T−Cytotoxic
CD8+T−PreExhaustion
CD8+T−Exhaustion
CD8+T−Memory
CD8+T−Effector
CD8+T−Naive
TRegs
CD4+Th17
TRegs−Resident
CD4+naive
CD4+Th1
pDC
cDC2
cDC1
Endothelial
Hepatocyte
MonocytesPatrolling
Monocytes
Macrophages−Capsule
MonocytesProliferating
KupfferCells
Neutrophils−Mature
Neutrophils−IFN
NeutrophilsImmature
NK−Resident
NK

**Fig. S3**

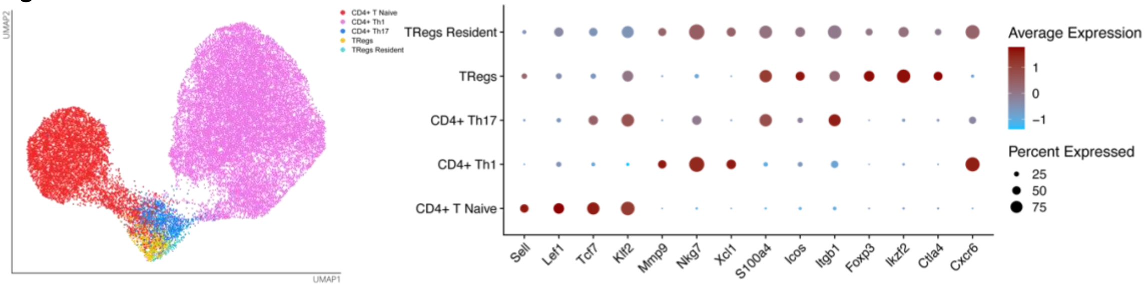
UMAP2
UMAP1
CD4+ T Naive
CD4+ Th1
CD4+ Th17
TRegs
TRegs Resident
TRegs Resident
TRegs
CD4+ Th17
CD4+ Th1
CD4+ T Naive
Sell
Lef1
Tcf7
Klf2
Mmp9
Nkg7
Xcl1
S100a4
Icos
Itgb1
Foxp3
Ikzf2
Ctla4
Cxcr6
Average Expression
1
0
−1
Percent Expressed
25
50
75

Fig. S4

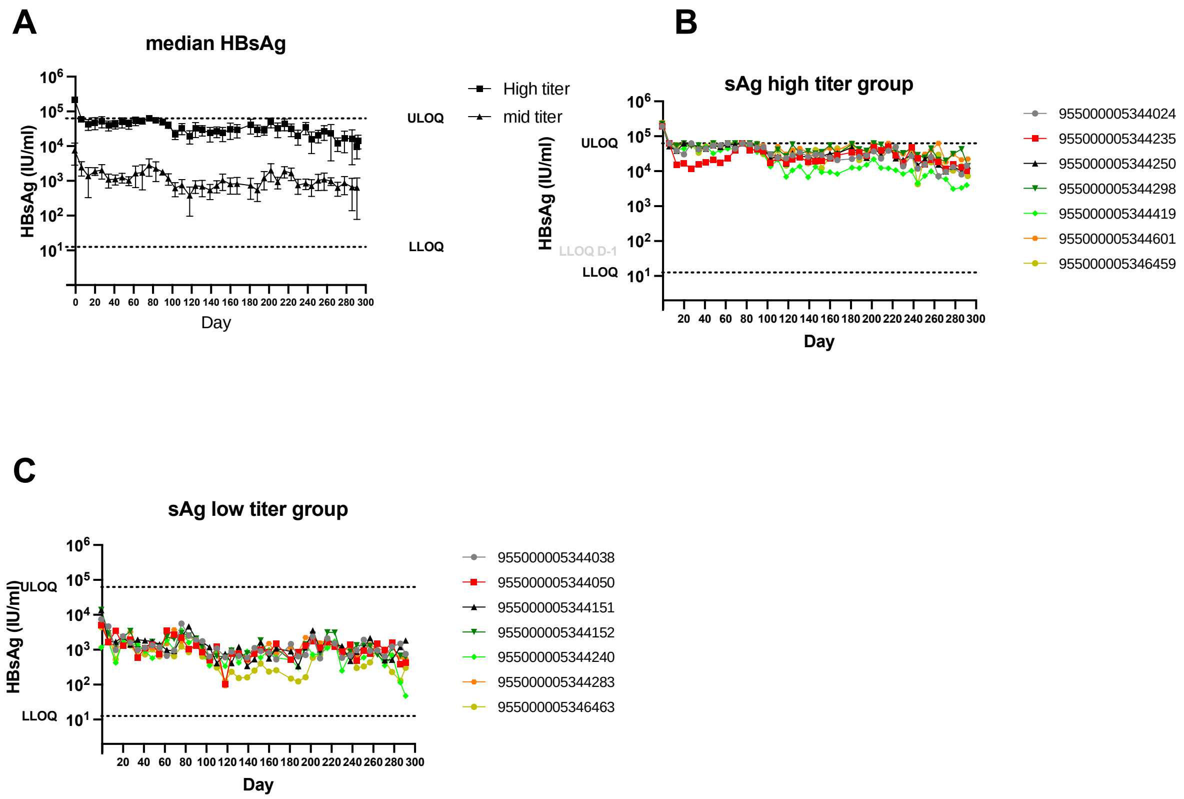
A
median HBsAg
High titer
mid titer
ULOQ
LLOQ
HBsAg (IU/ml)
Day
B
sAg high titer group
955000005344024
955000005344235
955000005344250
955000005344298
955000005344419
955000005344601
955000005346459
ULOQ
LLOQ D-1
LLOQ
HBsAg (IU/ml)
Day
C
sAg low titer group
955000005344038
955000005344050
955000005344151
955000005344152
955000005344240
955000005344283
955000005346463
ULOQ
LLOQ
HBsAg (IU/ml)
Day

**Fig. S5**

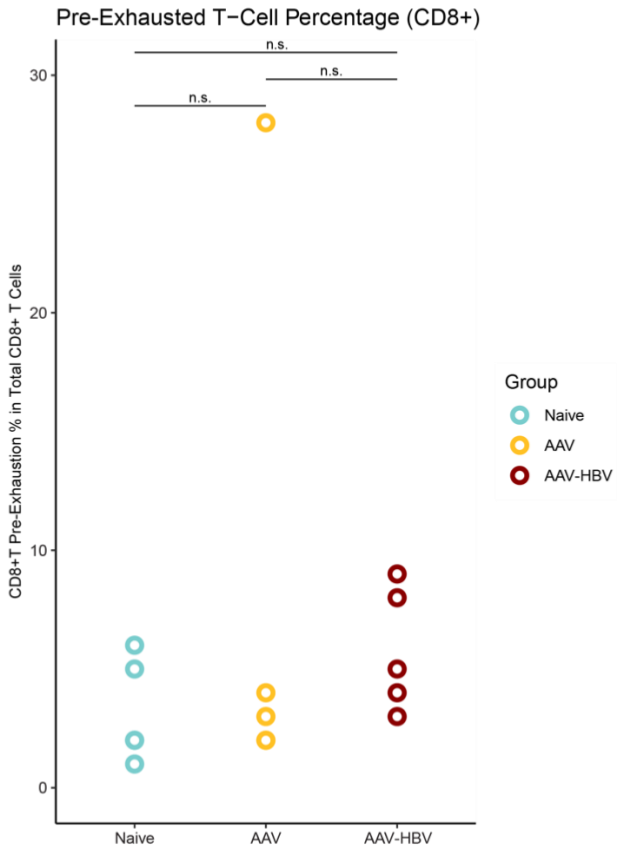

Pre-Exhausted T−Cell Percentage (CD8+)
n.s.
n.s.
n.s.
30
20
10
0
CD8+T Pre-Exhaustion % in Total CD8+ T Cells
Naive
AAV
AAV-HBV
Group
Naive
AAV
AAV-HBV

**Fig. S6**

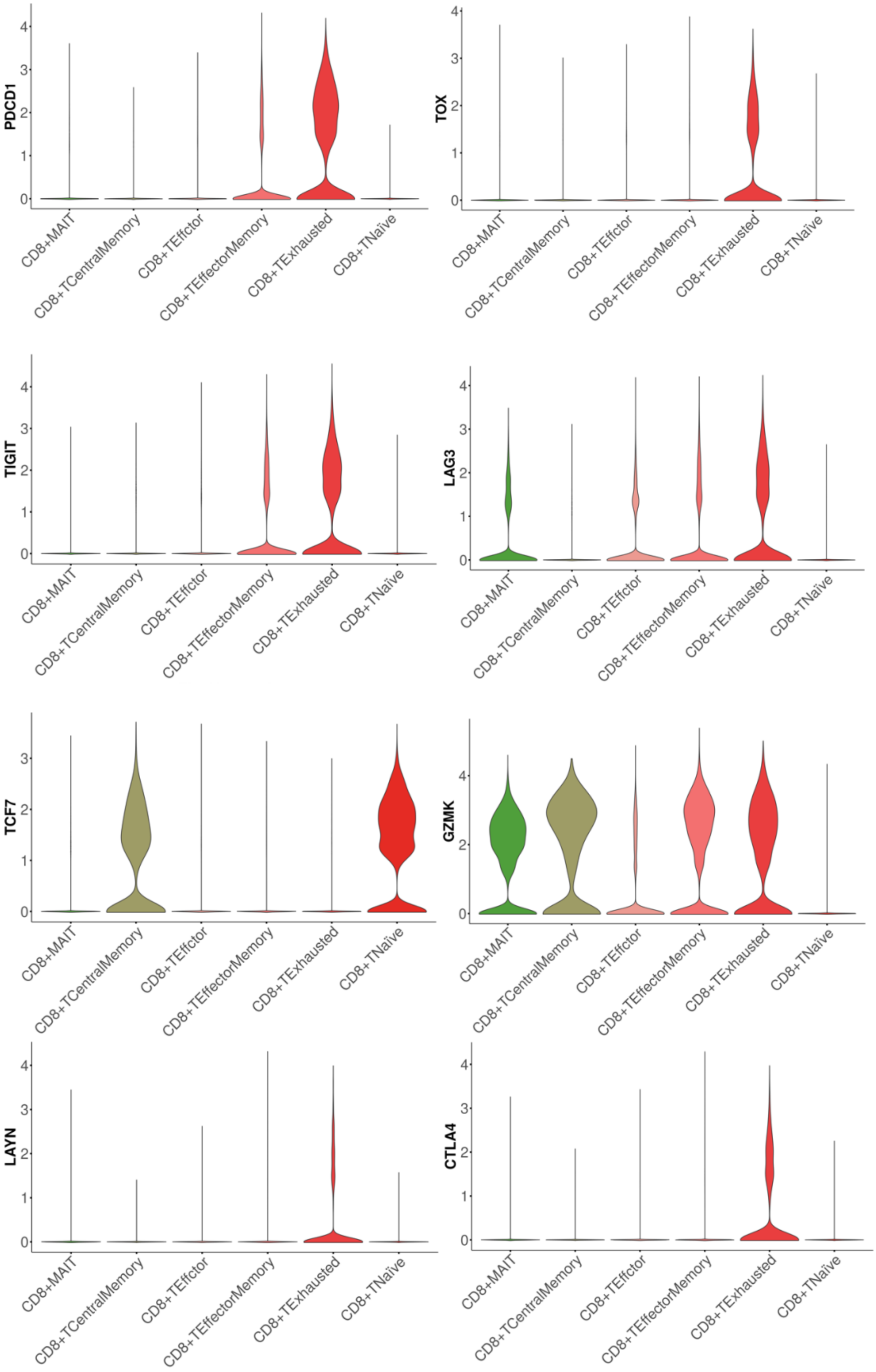
PDCD1
TOX
TIGIT
LAG3
TCF7
GZMK
LAYN
CTLA4
0
1
2
3
4
CD8+MAIT
CD8+TCentralMemory
CD8+TEffctor
CD8+TEffectorMemory
CD8+TExhausted
CD8+TNaïve

**SUPPLEMENTARY TABLES**

**Supplementary Table 1 – Cell Population Numbers and Frequencies**

| ***Cell Population*** | ***Cell Numbers*** | ***Frequency (%)*** |
|---|---|---|
| B-mature | 16121 | 20 |
| B-pre | 1628 | 2 |
| Basophils | 1165 | 1.5 |
| CD3+IFN+ | 644 | 0.8 |
| CD3+Proliferating | 900 | 1.1 |
| CD4+CD8 | 298 | 0.4 |
| CD4naive | 7247 | 9.1 |
| CD4Th1 | 20514 | 26 |
| CD4Th17 | 1015 | 1.3 |
| CD8T-Cytotoxic | 742 | 0.9 |
| CD8T-Effector | 711 | 0.9 |
| CD8T-Exhaustion | 785 | 1 |
| CD8T-Memory | 1949 | 2.4 |
| CD8T-Naive | 6245 | 7.8 |
| CD8T-PreExhaustion | 686 | 0.9 |
| cDC1 | 296 | 0.4 |
| cDC2 | 543 | 0.7 |
| Endothelial | 258 | 0.3 |
| Hepatocyte | 4248 | 5.3 |
| KupfferCells | 76 | 0.1 |
| Macrophages-Capsule | 328 | 0.4 |
| MAIT-Th17 | 943 | 1.2 |
| Monocytes | 2826 | 3.5 |
| MonocytesPatrolling | 634 | 0.8 |
| MonocytesProliferating | 405 | 0.5 |
| Neutrophils-IFN | 337 | 0.4 |
| Neutrophils-Mature | 1957 | 2.4 |
| NeutrophilsImmature | 324 | 0.4 |
| NK | 3108 | 3.9 |
| NK-Resident | 537 | 0.7 |
| NK-T | 838 | 1 |
| pDC | 362 | 0.5 |
| PlasmaCell | 640 | 0.8 |
| TRegs | 486 | 0.6 |
| TRegs-Resident | 174 | 0.2 |

**Supplementary table 2 – Marker genes for cell population typing**

| ***Cell Type*** | ***Marker genes*** |
|---|---|
| CD8 naïve | *Sell, Ccr7, Lef1, Tcf7, Klf2* |
| CD8 effector | *Gzma, Gzmb, Ccl5, Cx3cr1, Klrg1, S1pr5* |
| CD8 memory | *Sell, Ccl5, Ccr7, Eomes, Tcf7, Ly6c2* |
| CD8 cytotoxic | *Gzmb, Ccl5, Ccl4, Il7r, Gzma, Ccl3, Cd160* |
| CD8 precursor exhaustion | *Gzmk, Eomes, S100a6, Tox, Tigit, Klf2, Tcf7* |
| CD8 terminal exhaustion | *Gzmk, Lag3, Tox, Tigit, Pdcd1, Eomes, Nr4a2, Il10ra* |
| CD4 naïve | *Ccr7, Tcf7, Lef1, Sell, S1pr1, Ly6c1* |
| TRegs | *Foxp3, Ikzf2, Ctla4, Tigit, Capg, Tnfrsf4* |
| Tregs resident | *Cxcr6, Foxp3, Ikzf2, Ctla4, Tigit, Capg, Tnfrsf4* |
| Th1 | *Cxcr6, Ly6a, Il2rb, Klrb1c, Mmp9* |
| Th17 | *S100a4, Icos, Vim, Capg, Itgb1* |
| Cycling T-cell | *Top2a, Tuba1b, Mki67, Stmn1* |
| IFN stimulated T-cell | *Isg15, Ifit1, Ifit3, Cxcl10, Isg20, Rsad2, Irf7* |
| MAIT Th17 | *S100a4, Il7r, Tmem176a/b, Ramp1, Capg, Rorc, Pxdc1* |
| NK-T | *Fcer1g, Ly6c2, Klra7, Trdc, Ccl5, Anxa2, Cd7* |
| pre-B | *Iglc1, Cd24a, Sox4, Rgs2, Cd93* |
| Mature B | *Cd19, Cd79a, Cd79b, Ms4a1, Ly6d* |
| Plasma Cell | *Jchain, Igha, Igkc, Iglc1* |
| cDC1 | *Clec9a, Cd24a, Wdfy4, Id2, Ppt1* |
| cDC2 | *Cd209a, Cd7, Klrd1, Lgals1* |
| pDC | *Bst2m Siglech, Ly6d, Cox6a2* |
| Endothelial | *Prtn3, H2afy, Sox4* |
| Hepatocyte | *Fabp1, Car3, Apoe, Alb* |
| Monocytes | *Lyz2, Chil3, Hp, F13a1, Gn1* |
| Macrophages Capsule | *C1qa, C1qb, C1qc, Cd81, Apoe* |
| Monocytes Patrolling | *Ace, Ear2, Eno3, Gngt2* |
| Kupffer Cells | *C1qa, Vsig4, Cd5l, Folr2, Timd4, Clec4f* |
| Neutrophils Mature | *Cxcr2, Fcgr3, Csf3r, S100a8* |
| Neutrophils IFN | *Cxcr2, Rsad2, Isg15, Oasl1, Ifit1, S100a8* |
| Neutrophils Immature | *Ltf, Padi4, Lcn2, Camp, Mmp8, Ly6g, S100a8* |
| Basophils | *Cd63, Gata2, Csf1, Fcer1a, Cpa3, Hdc* |

**Supplementary table 3 –Statistics of frequencies between AAV-HBV high titer mice and AAV-control mice in all cell types.**

| Cell Type | group1 | group2 | n1 | n2 | statistic | p | p.adj | p.signif | p.adj.signif |
|---|---|---|---|---|---|---|---|---|---|
| B-cell | AAV | AAV-HBV | 5 | 5 | 11 | 0.841 | 1 | ns | ns |
| Basophils | AAV | AAV-HBV | 5 | 5 | 10 | 0.69 | 1 | ns | ns |
| CD3+ | AAV | AAV-HBV | 5 | 5 | 12 | 1 | 1 | ns | ns |
| DC | AAV | AAV-HBV | 5 | 5 | 10 | 0.69 | 1 | ns | ns |
| Endothelial | AAV | AAV-HBV | 5 | 5 | 10 | 0.69 | 1 | ns | ns |
| Hepatocyte | AAV | AAV-HBV | 5 | 5 | 12 | 1 | 1 | ns | ns |
| Monocytes | AAV | AAV-HBV | 5 | 5 | 13 | 1 | 1 | ns | ns |
| Neutrophils | AAV | AAV-HBV | 5 | 5 | 21 | 0.095 | 0.857 | ns | ns |
| NK | AAV | AAV-HBV | 5 | 5 | 5 | 0.151 | 1 | ns | ns |

**Supplementary table 4 –Statistics of frequencies between AAV-HBV high titer mice and AAV-control mice in T-cells.**

| CellType | group1 | group2 | n1 | n2 | statistic | p | p.adj | p.signif | p.adj.signif |
|---|---|---|---|---|---|---|---|---|---|
| CD3+IFN+ | AAV | AAV-HBV | 5 | 5 | 3 | 0.0556 | 0.222 | ns | ns |
| CD3+Proliferating | AAV | AAV-HBV | 5 | 5 | 4 | 0.0952 | 0.381 | ns | ns |
| MAIT-Th17 | AAV | AAV-HBV | 5 | 5 | 18 | 0.31 | 1 | ns | ns |
| NK-T | AAV | AAV-HBV | 5 | 5 | 17 | 0.421 | 1 | ns | ns |
| **CellType** | **group1** | **group2** | **n1** | **n2** | **statistic** | **p** | **p.adj** | **p.adj.signif** | **y.position** |
| CD4+CD8+ | AAV | AAV-HBV | 5 | 5 | 10 | 0.69 | 0.69 | ns | 0.03 |
| CD4+Th1 | AAV | AAV-HBV | 5 | 5 | 15 | 0.69 | 0.69 | ns | 0.79 |
| Tregs Resident | AAV | AAV-HBV | 5 | 5 | 14 | 0.84 | 0.84 | ns | 0.02 |
| CD4+naive | AAV | AAV-HBV | 5 | 5 | 14 | 0.84 | 0.84 | ns | 0.37 |
| CD4+Th17 | AAV | AAV-HBV | 5 | 5 | 9 | 0.55 | 0.55 | ns | 0.09 |
| TRegs | AAV | AAV-HBV | 5 | 5 | 6 | 0.22 | 0.22 | ns | 0.07 |
| **CellType** | **group1** | **group2** | **n1** | **n2** | **statistic** | **p** | **p.adj** | **p.adj.signif** | **y.position** |
| CD8+memory | AAV | AAV-HBV | 5 | 5 | 12 | 1 | 1 | ns | 0.33 |
| CD8+effector | AAV | AAV-HBV | 5 | 5 | 12 | 1 | 1 | ns | 0.13 |
| CD8+cytotoxic | AAV | AAV-HBV | 5 | 5 | 11 | 0.84 | 0.84 | ns | 0.19 |
| CD8 precursor exhaustion | AAV | AAV-HBV | 5 | 5 | 7 | 0.31 | 0.31 | ns | 0.31 |

| | | | | | | | | | |
|---|---|---|---|---|---|---|---|---|---|
| CD8 terminal exhaustion | AAV | AAV-HBV | 5 | 5 | 0 | 0.01 | 0.01 | ** | 0.19 |
| CD8+naive | AAV | AAV-HBV | 5 | 5 | 19 | 0.22 | 0.22 | ns | 0.73 |

**Supplementary table 5 – Flow cytometry and single-cell descriptive statistics of frequencies in AAV-HBV high titer mice**

| | **Single-Cell Exhausted** | **FC Exhausted** | **Single-Cell Pre-Exhausted** | **FC Pre-Exhausted** |
|---|---|---|---|---|
| Mean | 13.98 | 4.588 | 6.18 | 1.673 |
| Std. Deviation | 2.161 | 1.749 | 2.406 | 1.887 |
| Std. Error of Mean | 0.9666 | 0.7141 | 1.076 | 1.089 |
